# Cellular and molecular dynamics in the lungs of neonatal and juvenile mice in response to *E. coli*

**DOI:** 10.1101/2022.09.21.508849

**Authors:** Sharon A. McGrath-Morrow, Jarrett Venezia, Roland Ndeh, Benjamin D. Singer, Raffaello Cimbro, Mark Soloski, Alan L. Scott

## Abstract

Bacterial pneumonias cause significantly higher morbidity and mortality in neonates compared to other age groups. To understand the immune mechanisms that underlie these age-related differences, we employed a mouse model of *E. coli* pneumonia to examine cellular and molecular dynamics in immune responsiveness in neonates (PND 3-5) and juveniles (PND 12-18) at 24, 48, and 72 hours. Cytokine gene expression from whole lung extracts was quantified using qRT-PCR. *E. coli* challenge resulted in rapid and significant increases in neutrophils, monocytes, and γδT cells and significant decreases in dendritic cells and alveolar macrophages for both neonates and juveniles. Juveniles had significant increases in interstitial macrophages and recruited monocytes that were not observed in neonatal lungs. Expression of IFNγ-responsive genes were positively correlated with the levels and dynamics of MHCII-expressing innate cells in neonatal and juvenile lungs. Several facets of the responses of wild-type neonates was recapitulated in juvenile MHCII^-/-^ juveniles. Employing a pre-clinical model of *E. coli* pneumonia, we identified significant differences in the early cellular and molecular dynamics in the lungs that likely contribute to the elevated susceptibility of neonates to bacterial pneumonia and could represent targets for intervention to improve respiratory outcomes and survivability of neonates.

## Introduction

During early infancy and childhood, protective effector and memory immune responses in the lungs evolve through cumulative exposure to microbial and environmental antigens (1, 2). Because it takes time to accrue sufficient exposure to microbial antigens, children in the youngest cohorts have a higher likelihood of infection-induced severe disease, respiratory impairment, and death from pneumonia (3–6). As infants age and accumulate antigenic experience, the lung’s immune responses mature and establish trained innate immunity as well as resident memory cells that allow for the timely, efficient, and robust response that can provide protection from severe disease.

This period of relative vulnerability to respiratory pathogens corresponds to a time of rapid alveolar growth; with most of this growth occurring during the first two years of life (7). Growth in the lung is accompanied by Th2-mediated tissue remodeling (8, 9) that can be easily disrupted by Th1 and Th17 inflammation induced by microbial challenges. An attenuated pathogen-induced immune response may be important to allow for ongoing lung growth by minimizing cell cycle growth arrest responses and detrimental fibrotic remodeling. As such, the modified pathogen immune responses in infants may be beneficial or harmful depending on degree of virulence of the pathogen challenge (10, 11). While there are reports on the composition and functional status of alveolar macrophages, conventional dendritic cells, invariant natural killer T cells, and CD4 T cells in the neonatal lung at steady state after antigen challenge (1, 2, 8, 12–14), the cellular dynamics induced by live microbial challenge in the neonatal and juvenile lungs have received limited attention. In this study we sought to understand age-related differences in the cellular and molecular responses during the acute phase of the response induced by challenge with live *Escherichia coli*, a pathogen commonly associated with human neonatal pneumonia (15–18). Employing a live-challenge model of *E.coli* pneumonia, we demonstrate that the cellular dynamics of innate cells in the lungs of neonatal animals markedly differs from that of juveniles. Of particular note was the attenuated interstitial macrophage and monocyte responses in the lungs from neonatal animals that was associated with diminished MHCII-mediated responsiveness. These results reveal important new age-related differences in pathogen-induced immune responsiveness in the neonatal lungs that have implications for understanding the pathogenesis, management, and treatment of severe pneumonia in neonates.

## Results

### Cellular dynamics in neonatal and juvenile lungs in response to E. coli challenge

Based on experience with the live *E. coli* challenge model in neonatal and juvenile C57BL/6 mice (19), the dose chosen to study the cellular and molecular dynamics during the first 72 hours after challenge was 2.4 × 10^6^ CFUs. At this challenge level we reproducibly achieved >90% survival in both age groups (**Supplemental Figure 2A**), while causing demonstrable inflammation and tissue damage (**Supplemental Figure 2B**).

Flow cytometry was used to define the changes in the CD45^+^ immune cell populations in the lungs of WT neonatal and juvenile animals at 24, 48, and 72 hours *post-E.-coli* challenge (PEC). In neonates, *E. coli* exposure resulted in a rapid increase in neutrophil trafficking to the lungs that persisted through 72 hours PEC (**Figure 1**). In contrast, while neutrophils also trafficked to the lungs of juvenile mice early, they dropped to control levels by 72 hours PEC. Although the proportions of CD4 T cells and B cells did not significantly change during the early response in neonates and juveniles, there was a trend in both age groups for an increase in γδT cells at the later time points (**Figure 1**).

**Figure 1.**
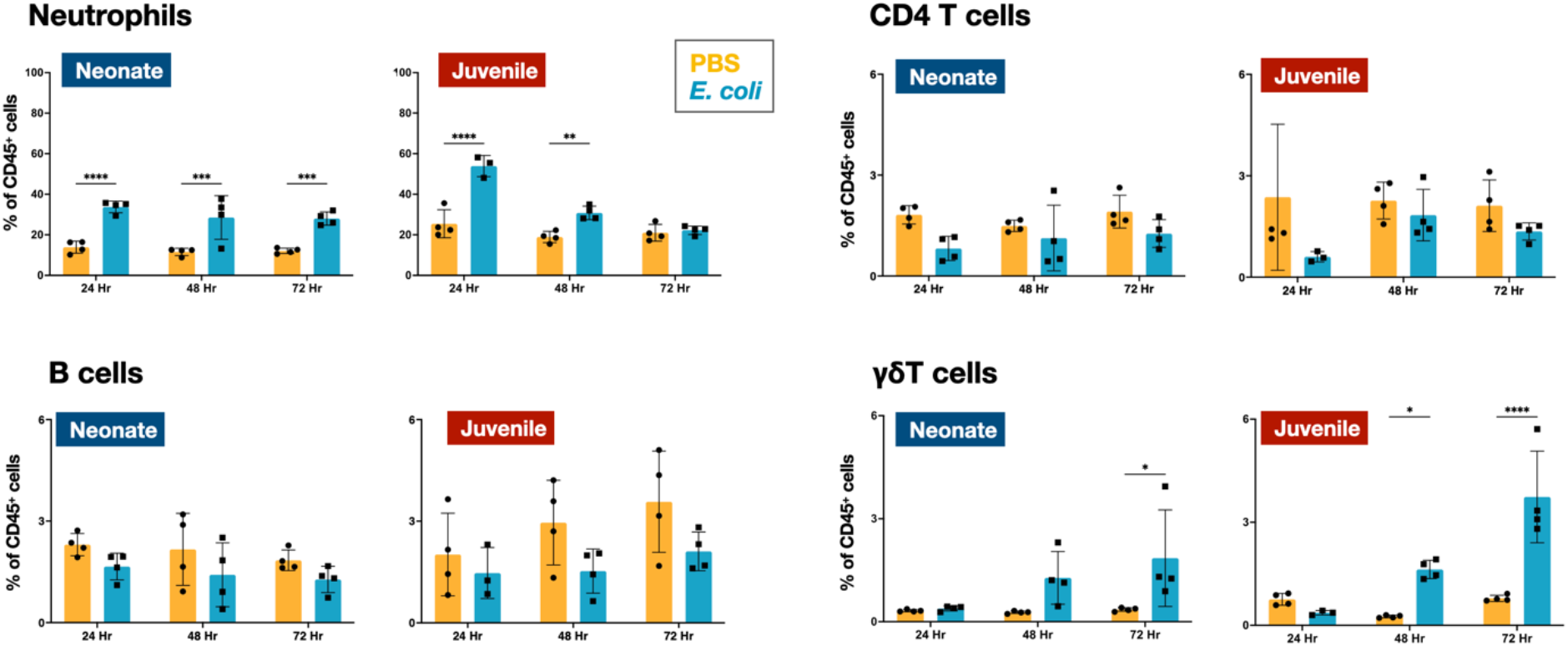
*E. coli*-induced changes in neutrophil and lymphocyte dynamics in neonatal and juvenile lungs. Changes in the dynamics of Ly6g^+^ neutrophils, CD4^+^ T cells, B220^+^ B cells, and γδ^+^ T cells in neonatal (PND 3-5) and juvenile (PND 12-18) lungs at 24, 48, or 72 hours PEC as assessed by flow cytometry (see **Supplemental Figure 1** for gating) and expressed as the percentage of total CD45^+^ cells. Statistical differences were determined using two-way ANOVA with multiple comparisons. Error bars represent standard deviation of the mean. ns - not significant; *p<0.05; **p<0.005; ***p=.0001; ****p<0.0001, (n = 4 group).

In the lungs of neonates, live *E. coli* challenge resulted in an early drop in the levels of dendritic cells (**Figure 2**) and alveolar macrophages (**Figures 2, 3A**) and these lower levels were sustained through 72 hours PEC. There was also a rapid drop in dendritic cells and alveolar macrophages in the lungs of juveniles, but in contrast to neonates, these cells recovered to control levels by 72 hours PEC, albeit with a distinct alteration in surface expression of F4/80 and SiglecF suggesting an altered activation status (**Figures 2, 3A**). While the lungs from *E. coli*-challenged juveniles displayed a sustained expansion of the interstitial macrophage compartment, there was no change to the constitutively low levels of this cell population in neonatal lungs (**Figures 2, 3B**). The dynamic changes in the levels of

**Figure 2.**
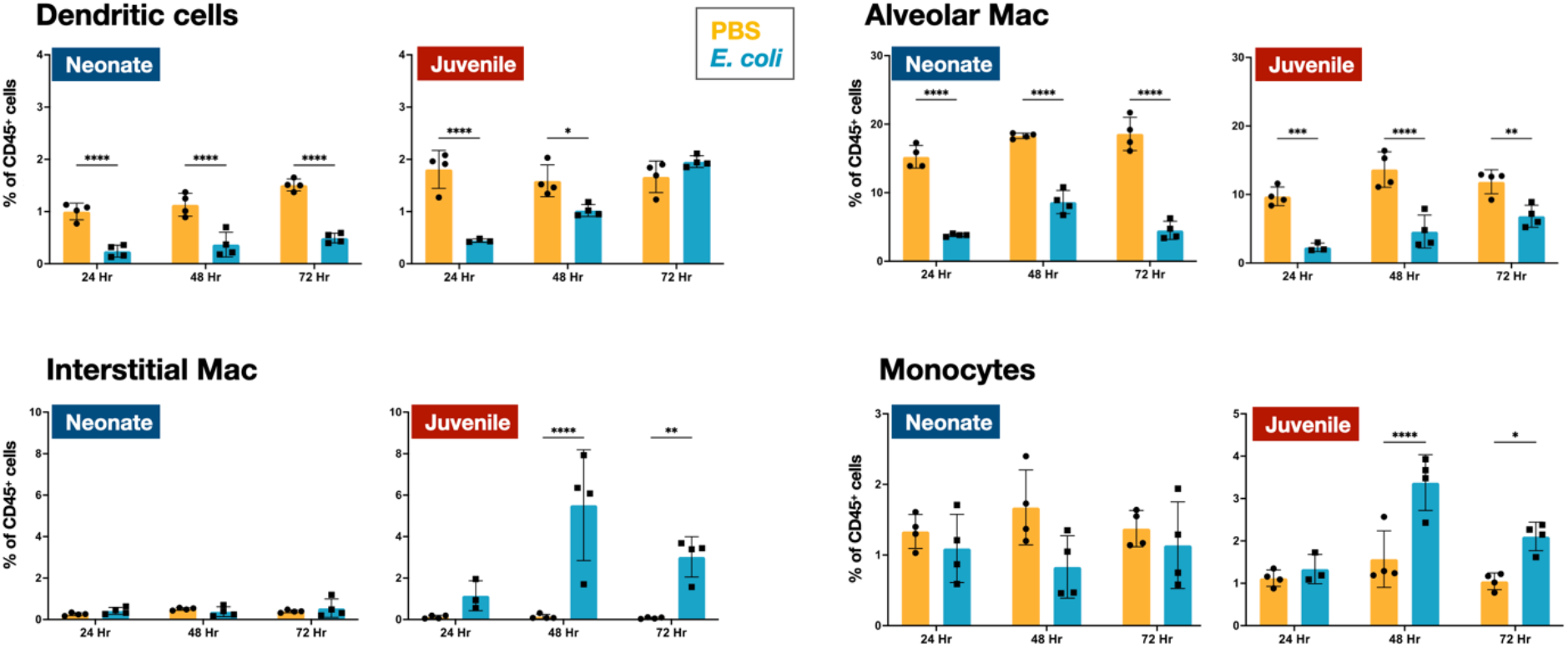
*E. coli*-induced changes in mononuclear cell dynamics in neonatal and juvenile lungs. Changes in the dynamics of CD11c^+^SiglecF^-^F4/80^-^MHCII^hi^ dendritic cells, CD11c^+^F4/80^+^SiglecF^+^ alveolar macrophages, CD11b^+^F4/80^+^SiglecF^-^ interstitial macrophages, and CD11b^+^Ly6c^+^MHCII^+^ monocytes in neonatal (PND 3-5) and juvenile (PND 12-18) lungs at 24, 48, or 72 hours PEC as assessed by flow cytometry (see **Supplemental Figure 1** for gating) and expressed as the percentage of total CD45^+^ cells. Statistical differences were determined using two-way ANOVA with multiple comparisons. Error bars represent standard deviation of the mean. ns - not significant; *p<0.05; **p<0.005; ***p=.0001; ****p<0.0001, (n = 3 or 4/group).

**Figure 3.**
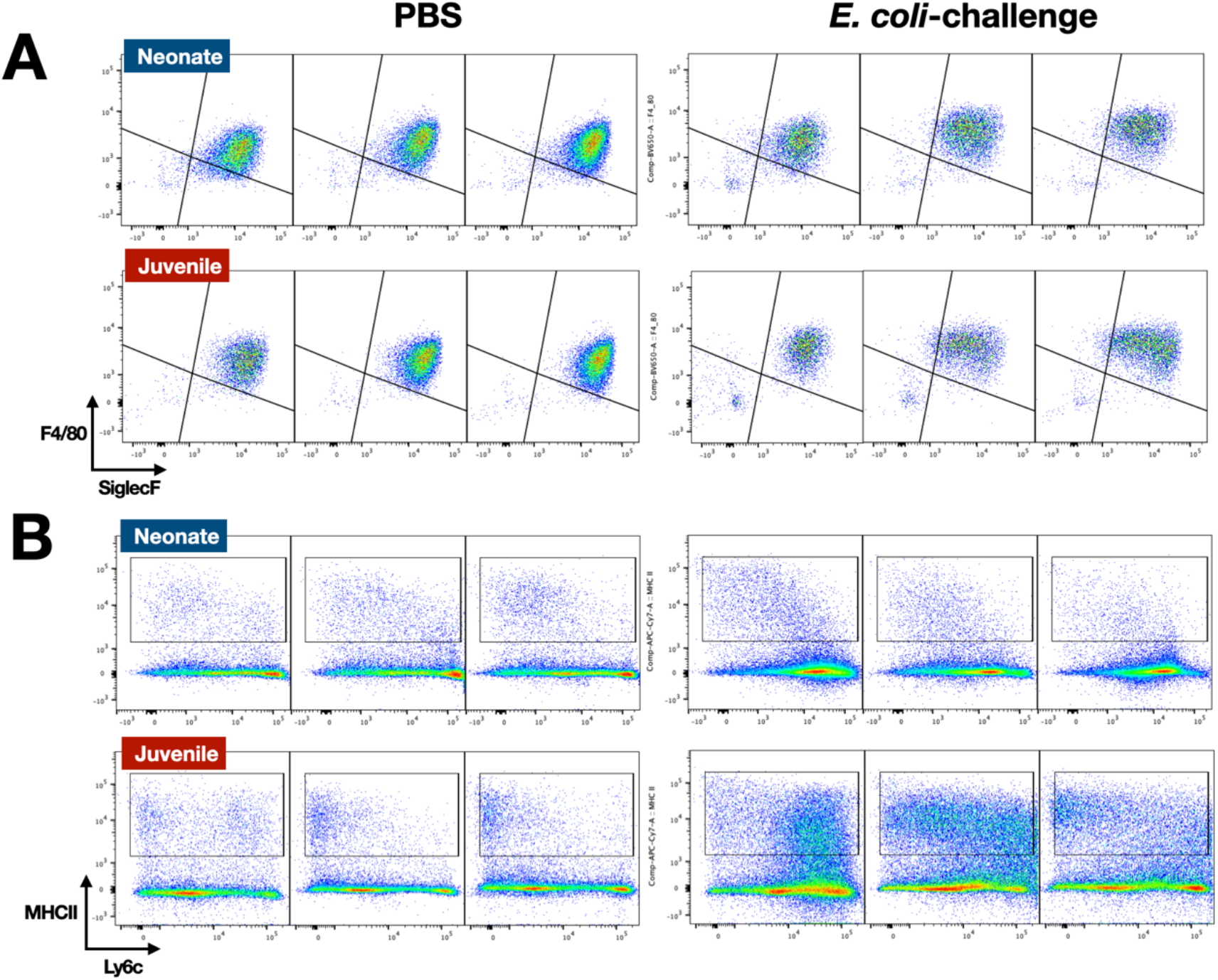
*E. coli*-induced changes in the surface phenotypes of alveolar macrophage and interstitial macrophage compartments. A. A representative flow cytometry analysis of F4/80^+^SiglecF^+^ alveolar macrophages from *E. coli*-challenged or PBS-treated neonatal (PND 3-5) and juvenile (PND 12-18) lungs at 24, 48, and 72 hours (see **Supplemental Figure 1** for gating).
B. A representative flow cytometry analysis of the CD11b^+^ MHCII^+^ Ly6c^variable^ interstitial macrophage compartment from *E. coli*-challenged or PBS-treated neonatal and juvenile lungs at 24, 48, and 72 hours (see **Supplemental Figure 1** for gating).

Ly6c expression on the surface of CD11b^+^MHCII+ cells, a heterogenous population that included interstitial macrophages (**Figure 3B**) suggest that recruited Ly6c^hi^ monocytes contributed to the expansion of this compartment in juveniles in response to *E. coli* challenge as reported by others (20, 21). This putative recruited monocyte-to-interstitial macrophage transition after *E. coli* challenge was not observed in the neonatal lungs. Although there was a significant increase in the proportion of monocytes in the juvenile lungs by 48 hours PEC, the proportion of monocytes in the neonatal lungs did not significantly change (**Figure 2**). This analysis identified differences in granulocyte, lymphoid and mononuclear cell dynamics between neonates and juveniles that might contribute to the enhanced sensitivity of neonatal animals to live *E. coli* challenge (19). Of particular note were the disparate temporal dynamics for dendritic cells, alveolar macrophages, interstitial macrophages, and monocytes, all cells that play key roles in innate immunity, tissue repair and initiating MHCII-mediated adaptive immune responses.

### Cellular dynamics in the lungs of neonatal and juvenile MHCII-/-mice to E. coli challenge

The differences in the dynamics of dendritic cells, alveolar macrophages, interstitial macrophages, and monocytes suggests that MHCII-mediated responses might account in part to the heightened morbidity and mortality in neonates. Indeed, previous work demonstrated that blocking MHCII during LPS challenged resulted in neonatal-like responses in the lungs of juvenile mice (22). With this in mind, we challenged MHCII^-/-^ animals to gain an insight into the influence of MHCII-mediated cellular and molecular responses in the lungs of neonatal and juvenile animals.

The MHCII^-/-^ neonates and juveniles had 80-85% survival at the 72 hour mark after the challenge with 2.4 × 10^6^ CFUs (**Supplemental Figure 2C**). In the lungs of MHCII^-/-^ neonates and juvenile there were notable levels of cellular infiltration in response to *E. coli*-challenge (**Supplemental Figure 2D**).

We examined the cellular immune dynamics in the lungs from neonatal and juvenile MHCII^-/-^ mice at 48 hours PEC; a timepoint chosen to represent the peak of lung inflammation induced by the *E. coli* challenge. The total CD45^+^ cell count significantly increased in both WT neonatal and WT juvenile lungs in response to *E. coli*-challenge, with a substantively larger growth in the juvenile lungs (**Figure 5**). In contrast, CD45^+^ cell dynamics was muted in the lungs from MHCII^-/-^ animals. As observed in WT animals, MHCII^-/-^ neonates and juveniles had an increase in the percentages and numbers of neutrophils at 48 hours PEC (**Figures 1, 4, 5**). Although γδT cells showed a proportional increase in the MHCII^-/-^ lungs (**Figure 4**), this was not reflected in an increase in γδT cell numbers as seen in the WT animals (**Figure 5**). CD4 T cells were largely absent from the MHCII^-/-^ lungs from both age groups. The proportions and numbers of B cells, dendritic cells, and monocytes found in MHCII^-/-^ neonatal and juvenile lungs at 48 hours PEC were similar to those found in WT animals (**Figures 1, 2, 4, 5**). Interestingly, compared to the response in WT lungs, the alveolar macrophage response to *E. coli* challenge remained blunted for both MHCII^-/-^ age groups at 48 hour PEC. Although the proportions of interstitial macrophages and monocytes were modestly higher in both neonatal and juvenile MHCII^-/-^ lungs (**Figure 4**), these proportional increases corresponded to significant increases in cell numbers (**Figure 5**). Taken together, these results suggest that MHCII signaling contributes to the early regulation of proliferation and/or activation of key innate cells in the neonatal lung environment.

**Figure 4.**
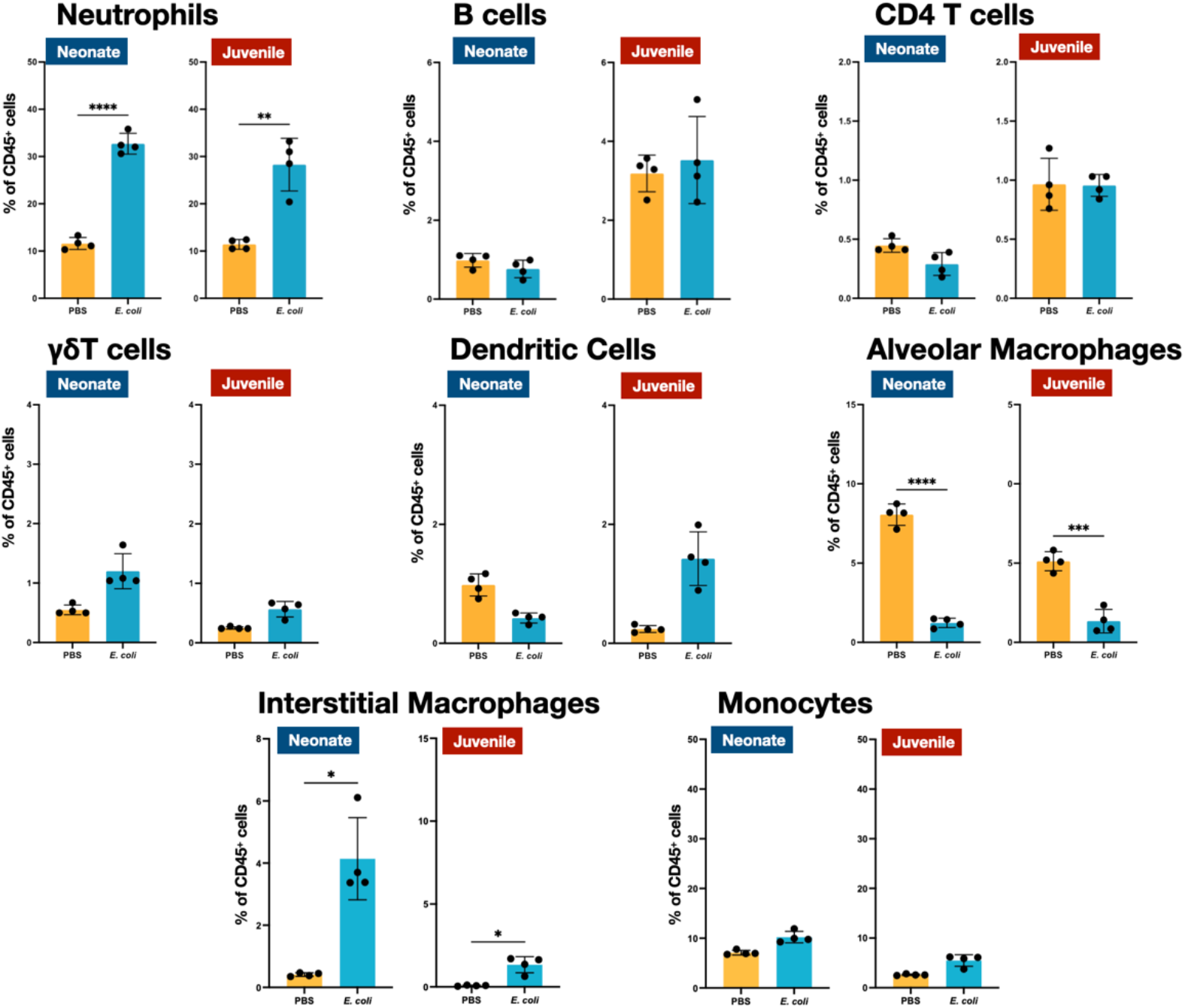
*E. coli*-induced changes in the proportions of neutrophils, lymphocytes and mononuclear cells in the lungs of MHCII-deficient neonates and juveniles at 48 hours post-challenge with *E. coli*. Changes in neutrophils, CD4 T cells, B cells, *γδ*T cells, dendritic cells, alveolar macrophages, interstitial macrophages and monocytes in neonatal (PND 3-5) and juvenile (PND 12-18) lungs at 48 hours PEC as assessed by flow cytometry (see **Supplemental Figure 1** for gating) and expressed as the percentage of total CD45^+^ cells. Statistical differences were determined using two-way ANOVA with multiple comparisons. Error bars represent standard deviation of the mean. ns - not significant; *p<0.05; **p<0.005; ***p=.0001; ****p<0.0001, (n = 3 or 4/group).

**Figure 5.**
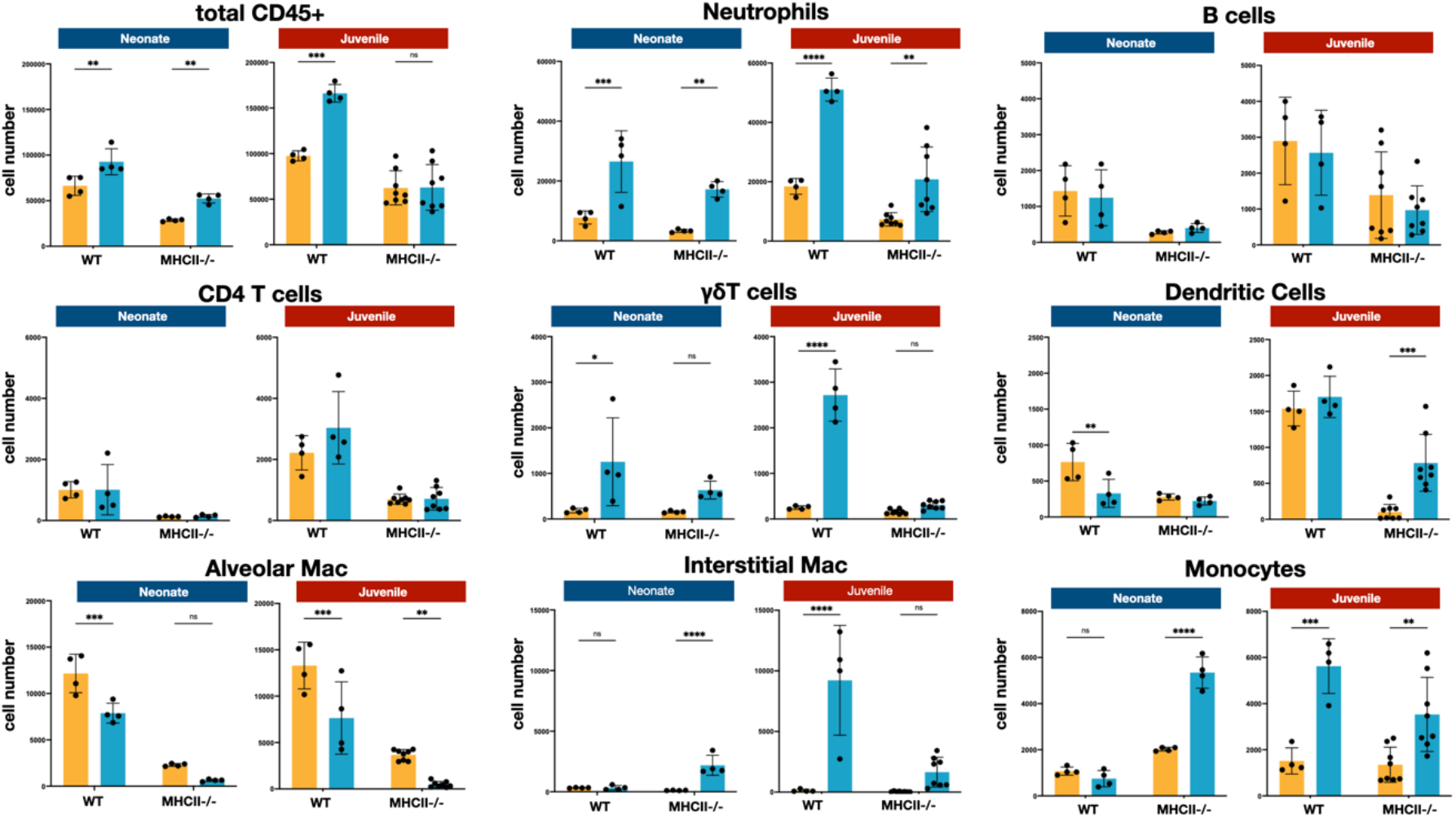
*E. coli*-induced changes in the numbers of neutrophils, lymphocytes and mononuclear cells in the lungs of WT and MHCII-deficient neonates and juveniles at 48 hours post-challenge with *E. coli*. Changes in the numbers of total CD45+ cells, neutrophils, CD4 T cells, B cells, *δδ*T cells, dendritic cells, alveolar macrophages, interstitial macrophages and monocytes in neonatal (PND 3-5) and juvenile (PND 12-18) lungs at 48 hours PEC as assessed by flow cytometry (see **Supplemental Figure 1** for gating). Statistical differences were determined using two-way ANOVA with Holm-Šídák post-hoc test for multiple comparisons. Error bars represent standard deviation of the mean. ns - not significant; *p<0.05; **p<0.005; ***p=.0001; ****p<0.0001, (n = 4 to 8/group).

### Transcriptional response in neonatal and juvenile lungs to E. coli challenge

The transcriptional dynamics of select cytokine and chemokine genes were measured from whole tissue extracts to obtain insight into the differences into the cell activation and trafficking in neonatal and juvenile lungs after exposure to live *E. coli*. The proinflammatory cytokines TNF and IL-6 are induced early downstream of signaling of pattern recognition receptors (23). While *tnf* and *Il6* transcription was increased in neonatal and juvenile lungs from both WT and MHC^-/-^ animals, WT juvenile lungs had heightened responsiveness at 24 and 48 hours PEC compared to the other groups (**Figure 6**). As predicted from the enhanced numbers of MHCII-positive cells (**Figure 2**), the WT juvenile lungs had the highest fold increase in *Ciita* expression at 24 and 48 hours PEC while the lungs from WT neonates and MHCII^-/-^ animals had minimal expression of lung *Ciita* at any time point (**Figure 6**). The *Ciita* transcription pattern supports our prior findings in which we found increased expression of MHCII on bronchoalveolar lavage-derived macrophages from juvenile but not from neonatal mice challenged with LPS (22). Studies have also demonstrated that a lack of pathogen-induced *Ciita* expression in the lungs of WT neonatal and MHCII^-/-^ mice is associated with lower numbers of IFNγ-producing CD4 T cells (24–26).

**Figure 6.**
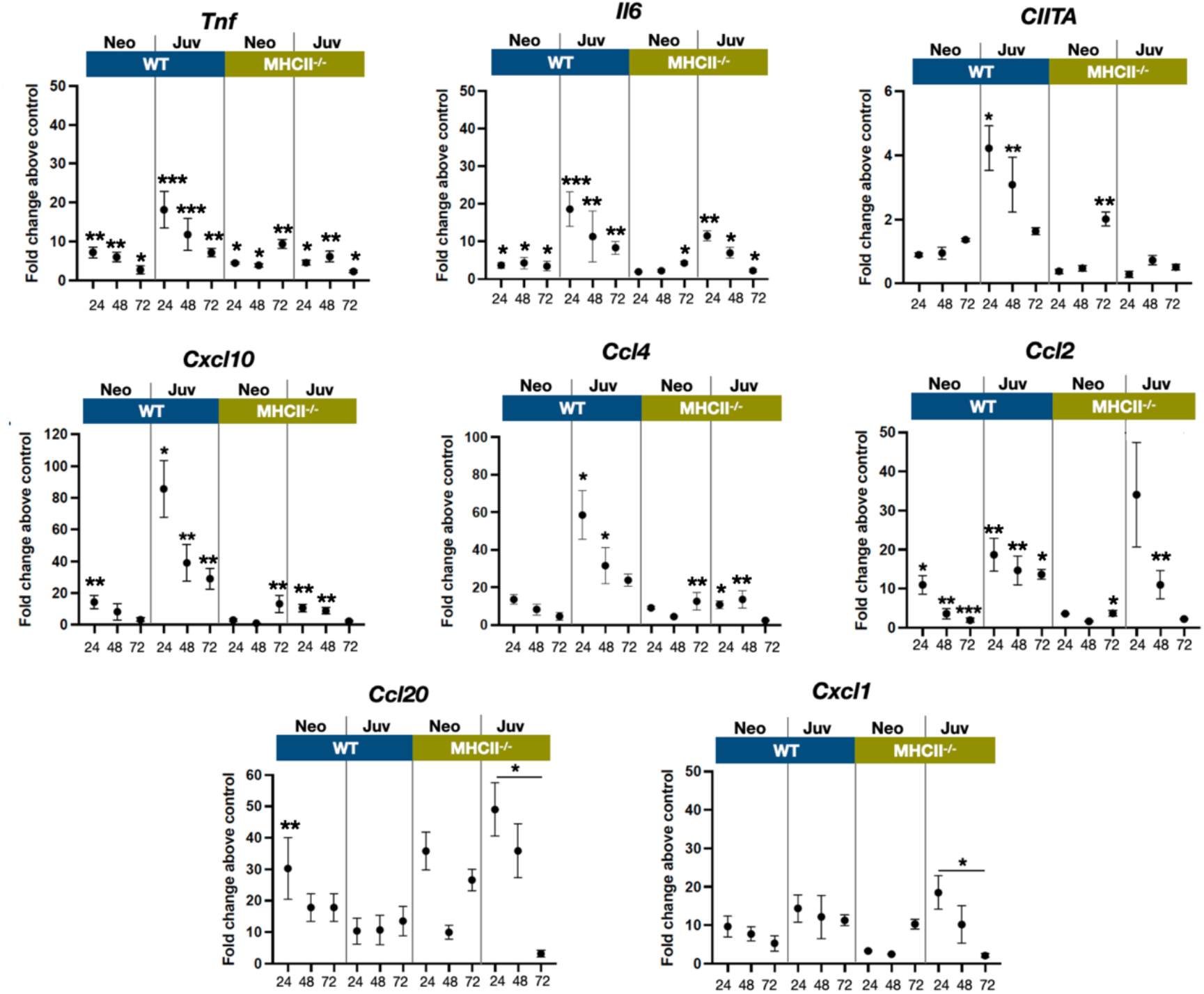
*E. coli*-induced changes in the expression dynamics of select regulatory, cytokine, and chemokine genes in WT and MHC^-/-^ neonatal and juvenile lungs. Quantitative RT-PCR analysis of total RNA isolated from the lungs of WT and MHCII-/-neonates (PND 3-5) and juveniles (PND 12-18) at 24, 48 and 72 hours post-*E. coli* challenge for changes in expression of class II major histocompatibility complex, transactivator (*Ciita*), tumor necrosis factor (*Tnf*), interleukin 6 (*Il6*), C-X-C motif chemokine ligand 10 (*Cxcl10*), C-C motif chemokine ligand 4 (*Ccl4*), C-C motif chemokine ligand 2 (*Ccl2*), C-C motif chemokine ligand 20 (*Ccl20*), and C-X-C motif chemokine ligand 1 (*Cxcl1*). Data presents as fold change over the value for the corresponding age/genetic background PBS controls. Points represent means of the values from 3-5 animals per group (error bars represent SEM). Statistical significance between groups were determined using one-way ANOVA with multiple comparisons. *p<0.01, **p<0.004, ***p<0.001, ****p<0.0001.

In support of the idea that low *Ciita* expression might be linked to low IFNγ production, the transcription of IFN-inducible genes *Cxcl10* were *Ccl4* were minimal in response to *E. coli* challenge in WT neonatal and MHCII^-/-^ animals (**Figure 6**). However, other IFN-inducible genes – *Ccl2, Ccl20* and *Cxcl1* – showed patterns of expression that suggested that additional factors contribute to the transcriptional regulation of these chemokine genes during the early stages of the innate response to *E. coli* challenge. *Cxcl1*, a Th17-associated chemokine involved in neutrophil migration and NFk*β* activation (27, 28), and *Ccl2*, a monocyte chemoattractant induced by the NFkß pathway (29), were also upregulated in MHCII^-/-^ juvenile lungs at 24 hours PEC (**Figure 6**). Interestingly, the fold changes in *Ccl20* transcription PEC were substantially lower in WT juvenile lungs compared to WT neonates or MHCII^-/-^ neonates and juveniles. In addition to its direct antibacterial properties (30), endothelial cell-derived CCL20 is reported to be chemotactic for CCR6^+^ lymphocytes and dendritic cells and to play a key roles in innate and adaptive inflammation at barrier surfaces (31). The early robust induction of *Ccl20* in WT neonate and MHCII^-/-^ lungs suggests a role for MHCII-mediated signaling in controlling the level of *Ccl20* transcription.

## Discussion

This study employed a murine model of bacterial pneumonia during infancy to examine age-related differences in the induction of lung immunity. The early responses of neonates and juveniles to live *E. coli* challenge were marked by substantial differences in the temporal dynamics for dendritic and mononuclear cells. Within 24 hours PEC there were significant reductions in both dendritic cells and alveolar macrophages that persisted in the neonatal lung but trended toward recovery to control levels by 72 hours PEC in the lungs from juveniles (**Figure 2**). Conventional dendritic cells are a prominent cell population during the first weeks of life in the mouse lungs that exhibit an activation profile that differs from adult-derived lung DCs by being strong inducers of Th2 immune responses (1). Colonization by the microbiota induces PD-L1 (programmed cell-death ligand 1) on neonatal lung dendritic cells, which endows them with the capacity to induce regulatory T cells (1). It is not clear if the persistent (neonates) or transient (juvenile) reductions observed here for dendritic cells and alveolar macrophages were due to cell death, emigration to local lymph nodes, or activation-associated changes in surface phenotype.

Lung macrophages are comprised of two major populations that occupy distinct anatomical compartments. Alveolar macrophages reside in the bronchoalveolar space and alveoli where these tissue resident cells function in regulating surfactant production, maintenance of epithelial barrier integrity, efferocytosis, and removal of inhaled particles (32). In addition, they have traditionally been designated as a first-line defense against air-borne pathogens, however, with the recent appreciation that recruited monocyte-derived macrophages are major contributors to the innate defense against microbial exposure, this aspect of alveolar macrophage function is under assessment (33, 34). The significantly less abundant interstitial macrophages are located in the interstitial spaces between the alveoli and the capillaries as well as in the interstitial environment surrounding conducting airways where they interact with stromal and other innate cells and play roles in inflammation (20, 21) and immunoregulation (35).

As noted, there was a significant reduction of alveolar macrophages at 24 hours PEC that persisted in neonatal lungs but was only transient in juvenile lungs (**Figure 2**). Acute loss of tissue resident macrophages at the early stages of inflammation, an occurrence referred to as ‘macrophage disappearance reaction’, has been described for peritoneal macrophages (36, 37) and in the lungs after LPS or CpG DNA challenge (32, 38). Interestingly, in these studies the transient loss of alveolar macrophages was accompanied by an expansion of interstitial macrophages (32, 38), similar to that seen in the lungs of WT juveniles challenged with live *E. coli* (**Figures 2, 5**).

A notable aspect of the cellular dynamics that distinguished the neonatal response from that of the juveniles was the striking differential in the numbers of interstitial macrophages (**Figures 2, 5**). Current evidence indicates that at steady state interstitial macrophages are maintained through slow replacement by monocytes that take up residency in the lungs (20, 21) and that a majority of the increases observed during inflammation is due to monocyte recruitment from the peripheral circulation (39). Thus, the interstitial macrophage levels are directly connected to the monocyte response, which was more robust in the lungs from juvenile animals. A majority of the increases in interstitial macrophages in juvenile lungs were made up of MHCII^hi^ cells (data not shown) which supports the idea that they were derived from recruited monocytes (20, 21) and could provide a cellular context for the increased *Ciita* expression observed in the whole lungs from juveniles. Given their role in the clearing of live bacterial challenges (40), the apparent lack of interstitial macrophage development in the neonatal lungs could be a contributing factor to the lower ability of neonates to clear a bacteria from the lung in a timely fashion and resolve inflammation (19).

The attenuated dynamics of MHCII-expressing mononuclear cells observed for neonates in this study likely contributes to the reported differences in MHCII-associated responsiveness between neonates and juveniles. Lee et al. (41) found that IFNγ-induced MHCII expression was attenuated in neonatal macrophages. LPS exposure significantly induced expression of MHCII in juvenile but not in neonatal peritoneal macrophages (22) and aspiration of LPS resulted in increased MHCII on and *Ciita* expression by bronchoalveolar lavage-derived mononuclear cells from juvenile but not from neonatal lungs (22). This reduced presence of MHCII-expressing cells might also have contributed to the reduced T cell responsiveness and diminished cytokine gene expression after live *E. coli* challenge. In general, there was an elevated transcription of *tnf, Il6*, and IFNγ-responsive genes in WT juvenile compared to WT neonatal and MHCII^-/-^ lungs (**Figure 6**). It is interesting to note that the responses in the lungs form juvenile MHCII^-/-^ animals at 48 hours PEC resembled the responses induced in the lungs from WT neonatal animals suggesting that restricted MHCII-associated signaling plays a key role in shaping the response in the neonatal lung. Studies carried out at the single cell level will likely be required to determine the relative contributions of MHCII^+^ mononuclear cell dynamics, the inherent propensity of CD4 T cells from juvenile mice to express Th1-associted cytokines upon *E. coli* challenge (10), or other factors in the differential responsiveness in neonatal and juvenile lungs.

Ideas and Speculation - It is possible that the attenuated inflammatory response of the neonate has some survival advantages. An overly robust immune response in the neonatal lung could disrupt essential growth dynamics during early postnatal lung development such that blunted responsiveness to pathogens during periods of rapid growth may be important in minimizing disruption to organogenesis while maintaining essential homeostatic and cellular processes. With a better understanding of the innate and adaptive responses in the neonatal lung environment it might be possible to devise interventions that transiently and strategically boost immune responsiveness in the neonate to improve respiratory outcomes to microbial challenge while minimizing the disruption to lung development. The findings reported here underscored differences in the cellular and molecular responses to live *E. coli* challenge that could contribute to the increased morbidity and mortality observed in neonates when challenged with a respiratory pathogen.

## Methods

### Mice

Timed pregnant C57BL/6NJ mice were obtained from NCI (Bethesda, MD). Animals were maintained on an AIN 76A diet and water *ad libitum* and housed at a temperature range of 20–23°C under 12-hour light/dark cycles. All experiments were conducted in accordance with the standards established by the United States Animal Welfare Acts, set forth in NIH guidelines and the Policy and Procedures Manual of the Johns Hopkins University Animal Care and Use Committee. MHCII-deficient mice on a B6 background were obtained from The Jackson Laboratory *(B6;129S2-H2^dlAb1-Ea^/JStock* No:003374 | MHC II^-^).

### Directly observed aspiration of *E. coli*

Pups were lightly sedated with isoflurane prior to *E. coli* (Seattle 1946, serotype O6, ATCC 25922) aspiration. Mice were given 2.4 × 10^6^ CFUs of *E. coli* in phosphate-buffered saline (PBS) or PBS alone. Forceps were used to gently extract the tongue, liquid was deposited in the pharynx and aspiration of fluid was directly visualized as previously described. (22) Neonatal mice (PND 3-5) received 10 μl of fluid and juvenile mice (PND 12-18) received 15 μl of fluid.

### Flow cytometry: Lung single cell suspension preparation

The lung was harvested, chopped to small pieces with a razor blade, and suspended in 1 ml of a buffer containing 1.0 mg DNAse 1 (Sigma) and 5.0 mg collagenase II (Worthington Biochem) in 1 ml RPMI. After incubating at 37°C for 30 minutes, an 18-gauge needle was used to further break up the lung tissue before passing the sample through a 70 μm cell strainer, into a 50 ml conical tube. PBS was added and the sample was centrifuged at 500×g for 5 minutes. The supernatant was discarded, and 1-2 ml of Ack lysing buffer was added and incubated at room temperature for 5 minutes. PBS was added to stop the reaction (3× to 5× the volume). The lung sample was then filtered through a 70 μm cell strainer into new 50 ml conical tube and spun down at 500xg for 5 minutes. FACS buffer was added (3 to 5 ml depending on the pellet size) and the cells were counted using a Biorad TC20 automated cell counter.

Cells were stained for viability using Live/Dead blue, UV (Invitrogen, L23105) and then with CD16/CD32 (BD Biosciences, 553142) to block Fc receptors. The cells were then stained with the following antibodies/fluorochromes: B220/AF488 (Biolegend, 103228), CD25/PE (BD, 561065), CD64/PEcy7 (Biolegend, 139314), CD11b/PE-CF594 (BD, 562317), CD4/BB700 (BD, 566408), CD11c/APC-R700 (BD, 565872), PD-1/APC-AF647 (Biolegend, 135209), MHC II/APC-Cy7 (Biolegend, 107627), SigF/BV421 (BD, 562681), Ly6G/BV510 (BD, 740157), Ly6C/BV605 (BD 128036), F4/80/BV650 (BD, 743282), CD3/BV711, (BD, 563123), ITCR gd/BV786/SB780 (Invitrogen, 78-5711-82), CD45/BUV395 (BD, 564279), and CD8/BUV737 (BD, 564297). Flow cytometry was performed using a BD Fortessa equipped with 5 lasers. The gating strategy used to identify lymphoid and myeloid cell populations is outlined in **Supplemental Figure 1**.

### Quantitative RT-PCR (qRT-PCR)

Total RNA was isolated from whole lung at 24, 48 and 72 hours post-*E. coli* challenge and analyzed as outlined previously (22). Reverse transcription was performed using total RNA and processed with the SuperScript first-strand synthesis system for RT-PCR according to the manufacturer’s protocol (Invitrogen). QRT-PCR was performed using the Applied Biosystems (Foster City, CA) TaqMan assay system, as previously described (22). Probes and primers were designed and synthesized by Applied Biosystems. The GADPH gene was used for an internal endogenous control. The following primer-probe sets were used: *Ccl2* (Mm00441242_m1), *Ccl4* (Mm00443111_m1), *Ccl20* (Mm01268754_m1), *Ciita* (Mm00482914_m1), *Cxcl1* (Mm04207460_m1), *Cxcl10* (Mm00445235_m1), *Il6* (Mm00446190_m1), and *Tnfα* (Mm00443258_m1).

### Statistical Analysis

Differences in measured variables between treated and control groups were determined using two-way ANOVA with or without Holm-Šídák post-hoc test for multiple comparisons. Comparison of survival curves were determined using log-rank (Mantel-Cox) test. Statistical significance was accepted at p<0.05. Error bars represent standard error of the mean.

---

**Supplemental Figure 1.**
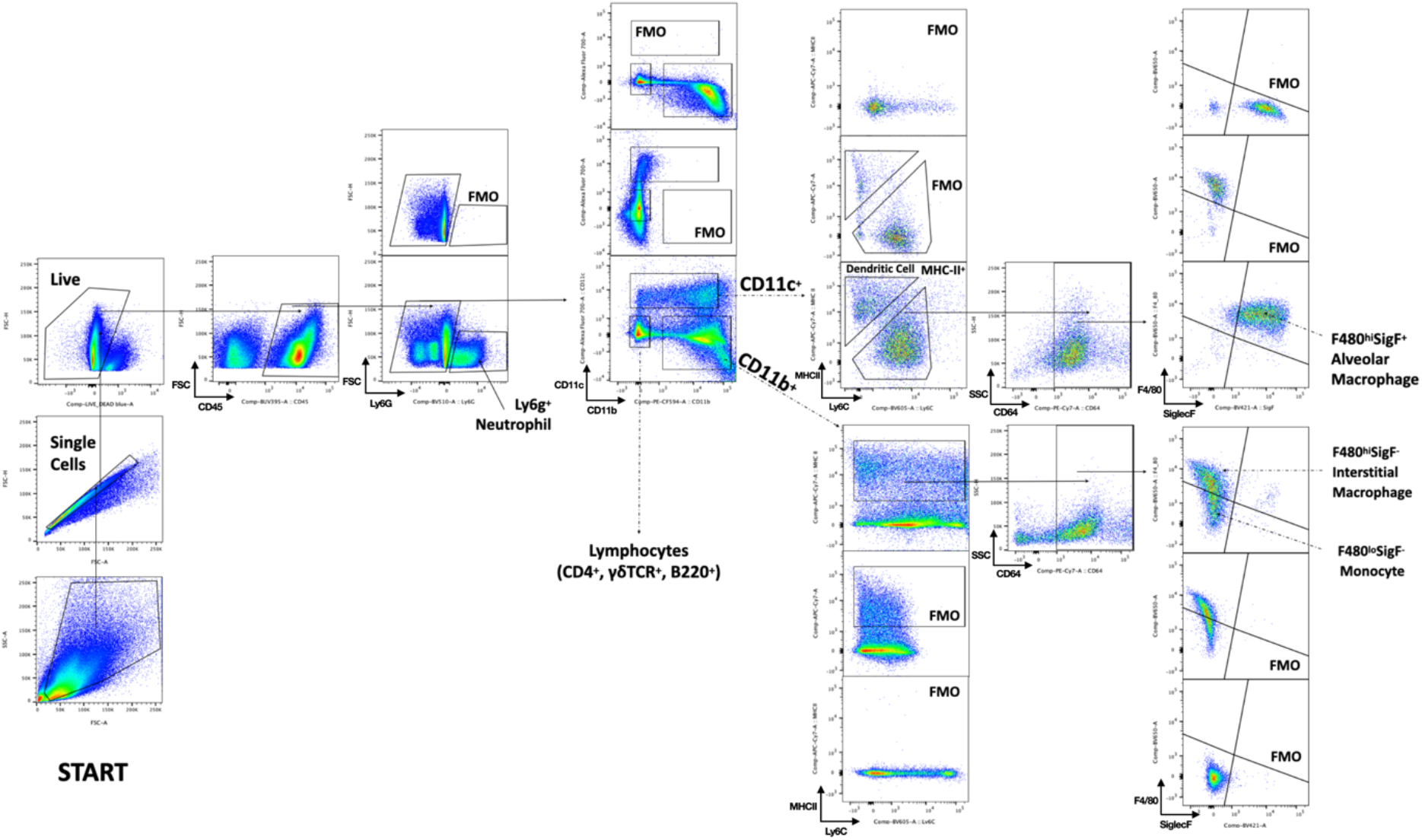
Gating Strategy. The gating strategy is illustrated using a juvenile (PND 12-18) 72-hour PBS-treated sample. Neutrophils were identified first among total CD45^+^ cells by Ly6g expression. Remaining cells were segregated by expression of CD11c or CD11b. Populations in the CD11c^+^compartment include MHC-II^hi^ dendritic cells and F480^+^ SiglecF^+^ alveolar macrophages. Cells within the CD11b^+^ compartment include two F480^lo^SiglecF^-^ monocyte populations with variable MHC-II expression, and MHC-II^hi^F480^hi^SiglecF^-^ interstitial macrophages.

**Supplemental Figure 2.**
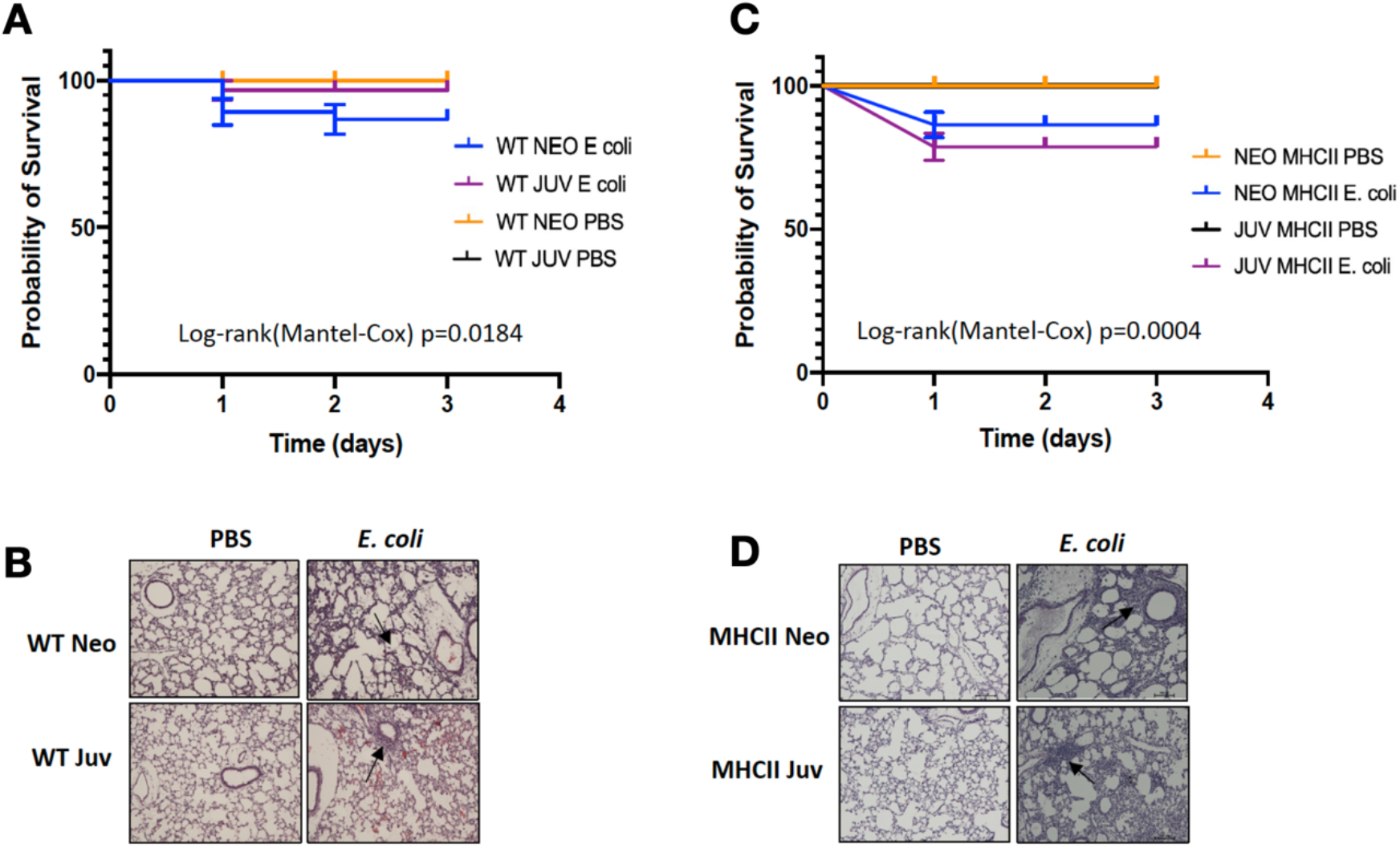
Survival analysis of WT and MHCII^-/-^ neonatal (PND 3-5) and juvenile (PND 12-18) animals after challenge with *E. coli*. A. Mortality of WT neonates and juveniles at 24, 48, and 72 hours post-*E. coli* challenge with 2.4 × 10^6^ CFUs or PBS treatment. Comparison of survival curves were determined using log-rank (Mantel-Cox) test. (n= 24-47/group)
B. Representative examples of the histological appearance of the lungs from WT neonates and juveniles at 48 hours post-*E. coli* challenge or control PBS-treatment (arrows point to inflammatory foci).
C. Mortality of MHCII^-/-^ neonates and juveniles at 24, 48, and 72 hours post-*E. coli* challenge with 2.4 × 10^6^ CFUs or PBS treatment. Comparison of survival curves were determined using log-rank (Mantel-Cox) test. (n=12-37/group)
D. Representative examples of the histological appearance of the lungs from MHCII^-/-^ neonates and juveniles at 48 hours post-*E. coli* challenge or control PBS-treatment (arrows point to inflammatory foci).

